# Determining the efficacy of hand sanitizers against virulent nosocomial infections

**DOI:** 10.1101/2020.10.12.335844

**Authors:** Jawad J Najdawi, Lester D Hubble, Alec J Scarborough, Matthew J Watz, Paul Watson

## Abstract

The goal of this study was to determine the effectiveness of 0.13% benzalkonium chloride (BAC) (Steirolotion™), 100% ethanol, and 70% ethyl alcohol (Purell™) hand sanitizer in subduing the growth of nosocomial bacteria – methicillin resistant Staphylococcus *aureus* (MRSA), vancomycin resistant Enterococcus (VRE), and Pseudomonas *aeruginosa* (P. *aeruginosa*) - when plated on culture media over an extended period of time. In addition, our objective was to quantify the length of time these hand sanitizer agents remained effective, and to extrapolate their efficacy in decreasing the transmission of hospital-acquired infections. 50 microliters of either BAC, 100% ethanol, or 70% ethyl alcohol hand gel sanitizer were pipetted onto Trypticase soy agar with 5% sheep blood plates that were cultured with either MRSA, VRE, or P. *aeruginosa*. The plates were then incubated at 37.0?. The zone of inhibition (ZOI) was measured daily for 5 days and additionally zones were noted whether or not regrowth recurred in areas where previous growth had initially been inhibited. BAC was found to be superior to both 100% ethanol and 70% ethyl alcohol in the inhibition of MRSA over all time points (p values < .05). BAC was found to be superior to 70% ethyl alcohol in the inhibition of VRE over all time points (p values < .05), but not statistically superior to 100% ethanol against the inhibition of VRE over any time points. BAC was found to be superior to 70% ethyl alcohol and 100% ethanol in the inhibition of pseudomonas over 72 and 24 hours, respectively (p values < .05). The results of this study demonstrate *in vitro* efficacy of BAC of preventing regrowth of common nosocomial bacteria over a prolonged period of time, especially when compared to ethyl alcohol. Further study is warranted to determine *in vivo* effectiveness of this formulation of BAC as well as the appropriate time frame of application for effectiveness against P. *aeruginosa*.

## Introduction

Hospital-acquired or healthcare-associated infections (HAIs) are the most common complication in hospitalized patients [1]. They occur with an estimated incidence of 4.5 HAIs per 100 hospital admissions, and amount to an additional burden of $35 to $45 billion dollars on the healthcare system [2]. They are responsible for significant hardship accounting for more than 90,000 deaths each year, putting HAIs among the top 5 leading causes of death in the United States [3-5]. Transmission of pathogens from healthcare staff serves as an important source of HAIs. Personal hygiene is a crucial aspect of reducing transmission, and hand washing, or sanitizing is required with every patient contact [6].

Both alcohol-based and alcohol-free hand sanitizers are available options when hand washing is not available or efficient. Alcohol-based sanitizers containing 60-95% alcohol are most often used in hospitals. Benzalkonium chloride (BAC) is the active ingredient contained in most alcohol-free hand sanitizer products available today. It has been theorized that BAC possesses an extended killing time of bacteria when the solution has dried compared to alcohol-based agents [7-10].

The goal of this study was to determine the duration of efficacy of .13% BAC, 70% ethyl alcohol, and 100% ethanol in decreasing methicillin-resistant Staphylococcus *aureus* (MRSA), vancomycin-resistant Enterococcus (VRE), and Pseudomonas *aeruginosa* (P. *aeruginosa*) colonization and regrowth over an extended period of time when plated on culture media.

## Materials and Methods

BD BBL™ Trypticase™ soy agar slants prepared media of nosocomial bacteria MRSA, VRE, and P. *aeruginosa* were grown on agar plates for 24 hours at 37 degrees Celsius and used to establish a reservoir.

A 0.5 McFarland standard solution was created for the MRSA, VRE, and P. *aeruginosa* bacteria strains, by using a calibrated inoculating loop to transfer bacteria from the incubated blood agar plates to a vial of saline with 0% absorbance until the absorbance of the vial solution was between 0.08 and 0.1%.

A bacteria lawn was created by using a cotton applicator to evenly distribute an aliquot of the 0.5 McFarland standard MRSA solution across the surface of Trypticase™ soy agar with 5% sheep blood plates. This process was repeated using the 0.5 Mcfarland standard for MRSA, VRE, and P. *aeruginosa* solutions until eight bacteria lawns of each solution were created.

50 microliters of 0.13% benzalkonium chloride (BAC) (SteirolotionTM, Germcure, Houma Louisiana), 100% ethanol (Sigma-Aldrich Inc., St. Louis, Missouri) and ethyl alcohol 70% (Purell™, Gojo, Akron, Ohio) solution were pipetted onto each of the eight 5% sheep blood agar plates. Reverse pipetting was used to ensure accurate amounts of viscous solution were pipetted onto the plates. Plates were left for one hour to dry.

The MRSA, VRE, and *P. aeruginosa* inoculated plates were incubated at 37 degrees Celsius overnight. They were all grown in aerobic conditions. The plates were removed from the incubator every 24 hours for a growth period of 120 hours to take photographs and quantitative measurements of the zone of inhibition (ZOI) of each antiseptic. Measurements were performed for a total of 120 hours for the MRSA, VRE, and P. aeruginosa plates.

One methodology was utilized to perform quantitative measurements of the ZOI for each antiseptic. The methodology used to perform digital measurements of the ZOI was the free internet software program, ImageJ. ImageJ utilizes the pixels of the digital photographs taken and the known standard diameters of the agar plates to quantitatively measure the ZOI. Four researchers made the digital measurements independently to increase validity of the measurements.

Statistical analysis was performed by measuring the difference in area of inhibition between ethyl alcohol, ethanol, and benzalkonium chloride for P. *aeruginosa*, VRE, and MRSA each. P-values were obtained using t-tests comparing each solution independently.

## Results

Ethyl alcohol and Ethanol showed significant regrowth of bacteria within 24 hours against P. *aeruginosa*, VRE, and MRSA (Figs 1-3). This regrowth of bacteria continued the full 5 days, or 120 hours, that the study was conducted. BAC showed regrowth of only P. *aeruginosa* after the initial 24-hour period had passed. For BAC, no regrowth was noted throughout the 120 hours in MRSA or VRE after the initial ZOI had been established (Figs 1 and 2, 4 and 5).

**Fig 1.**
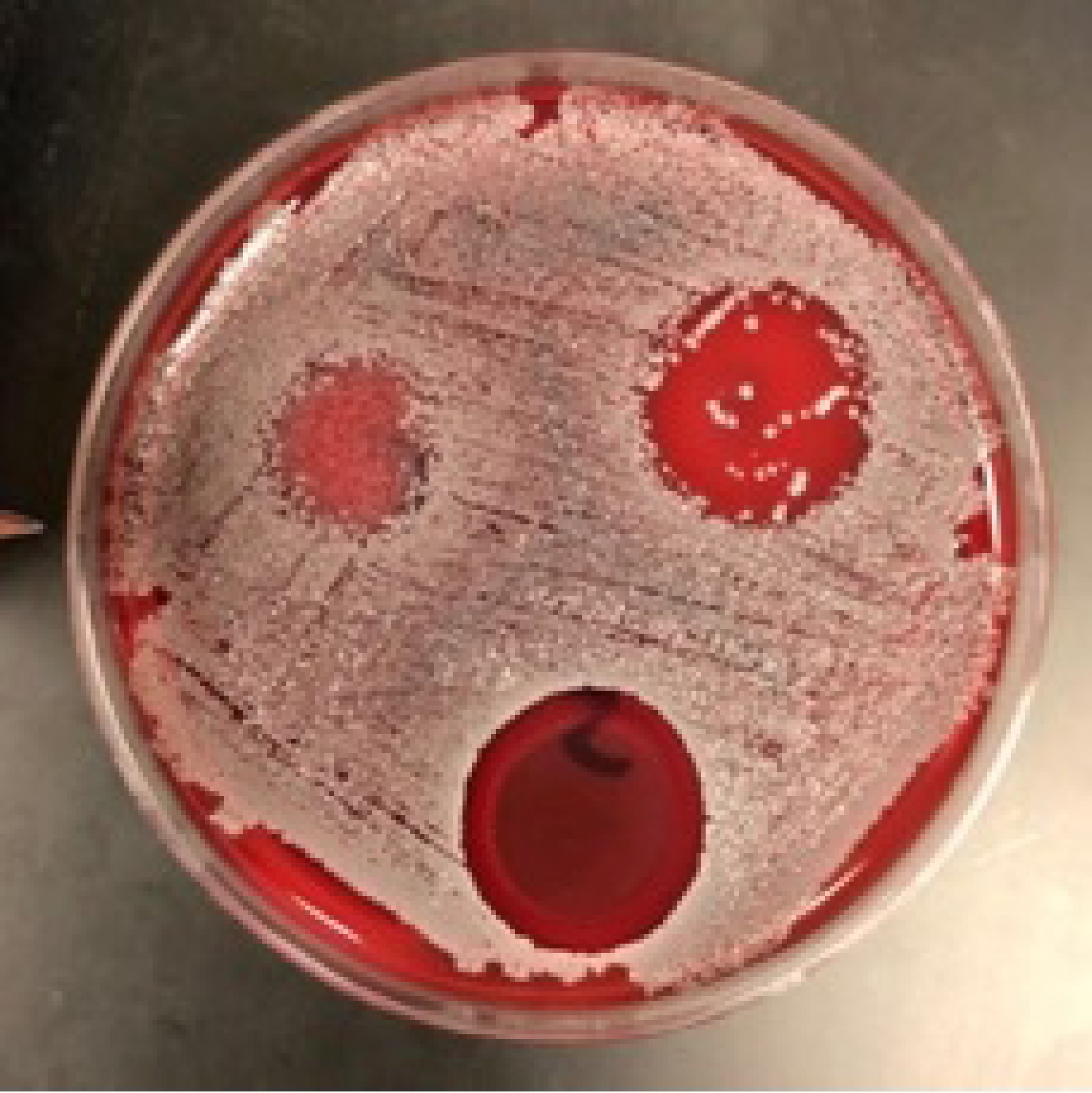
One MRSA plate after 24 hours in incubator. Ethyl alcohol (top left), Ethanol (top right), and BAC (bottom).

**Fig 2.**
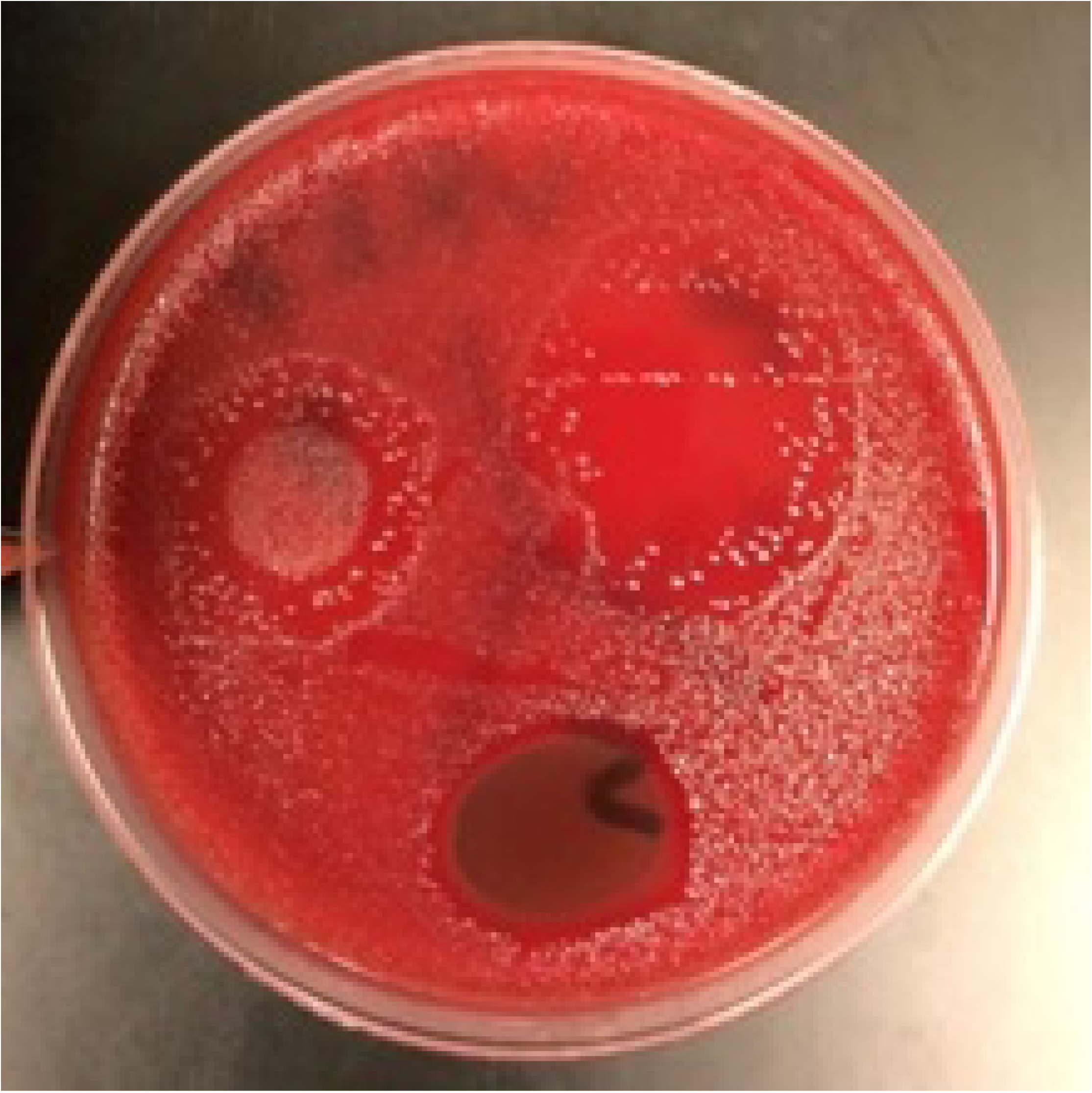
One VRE plate after 24 hours in incubator. Ethyl alcohol (top left), Ethanol (top right), and BAC (bottom).

**Fig 3.**
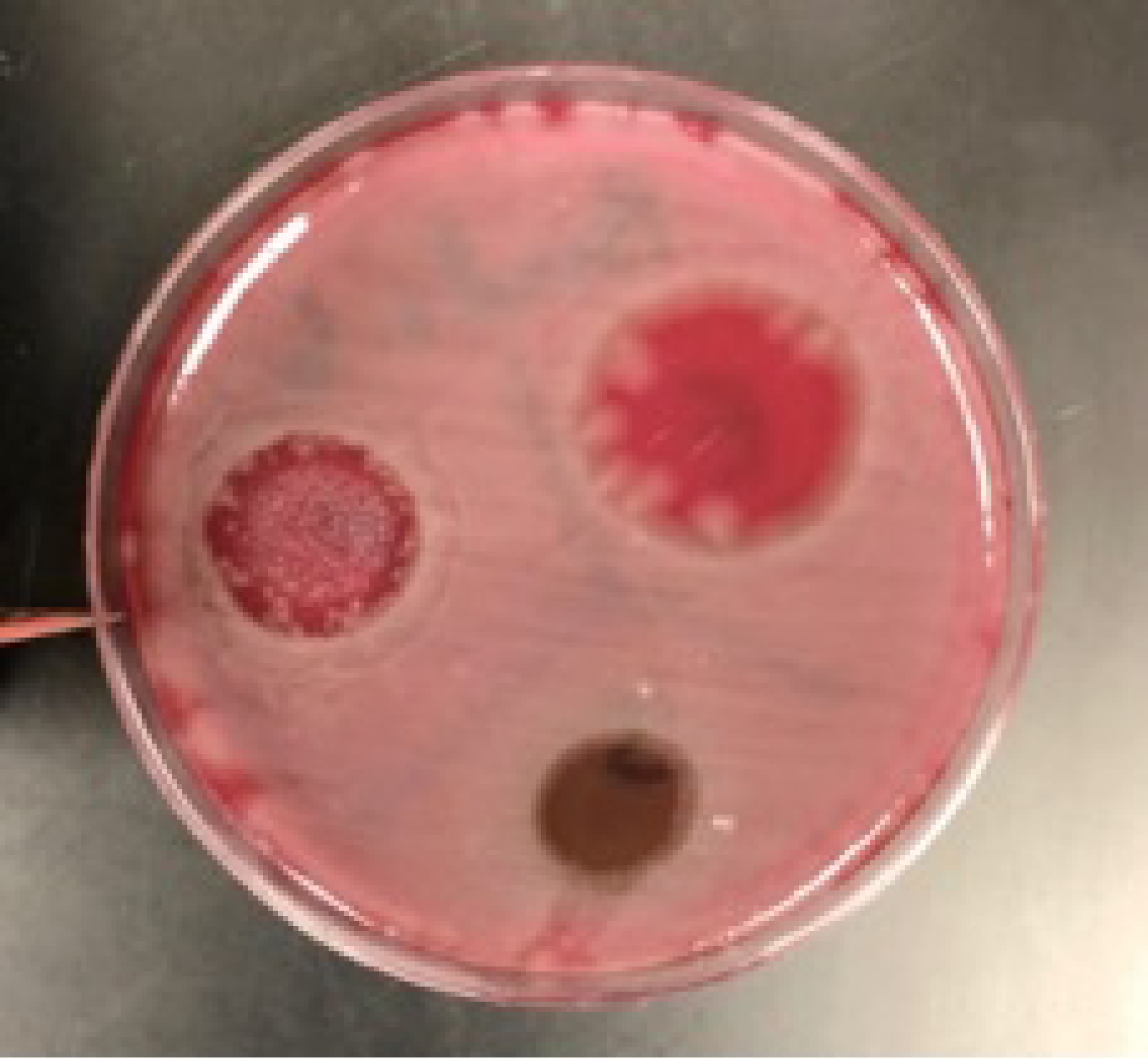
One P. *aeruginosa* plate after 24 hours in incubator. Ethyl alcohol (top left), Ethanol (top right), and BAC (bottom).

**Fig 4.**
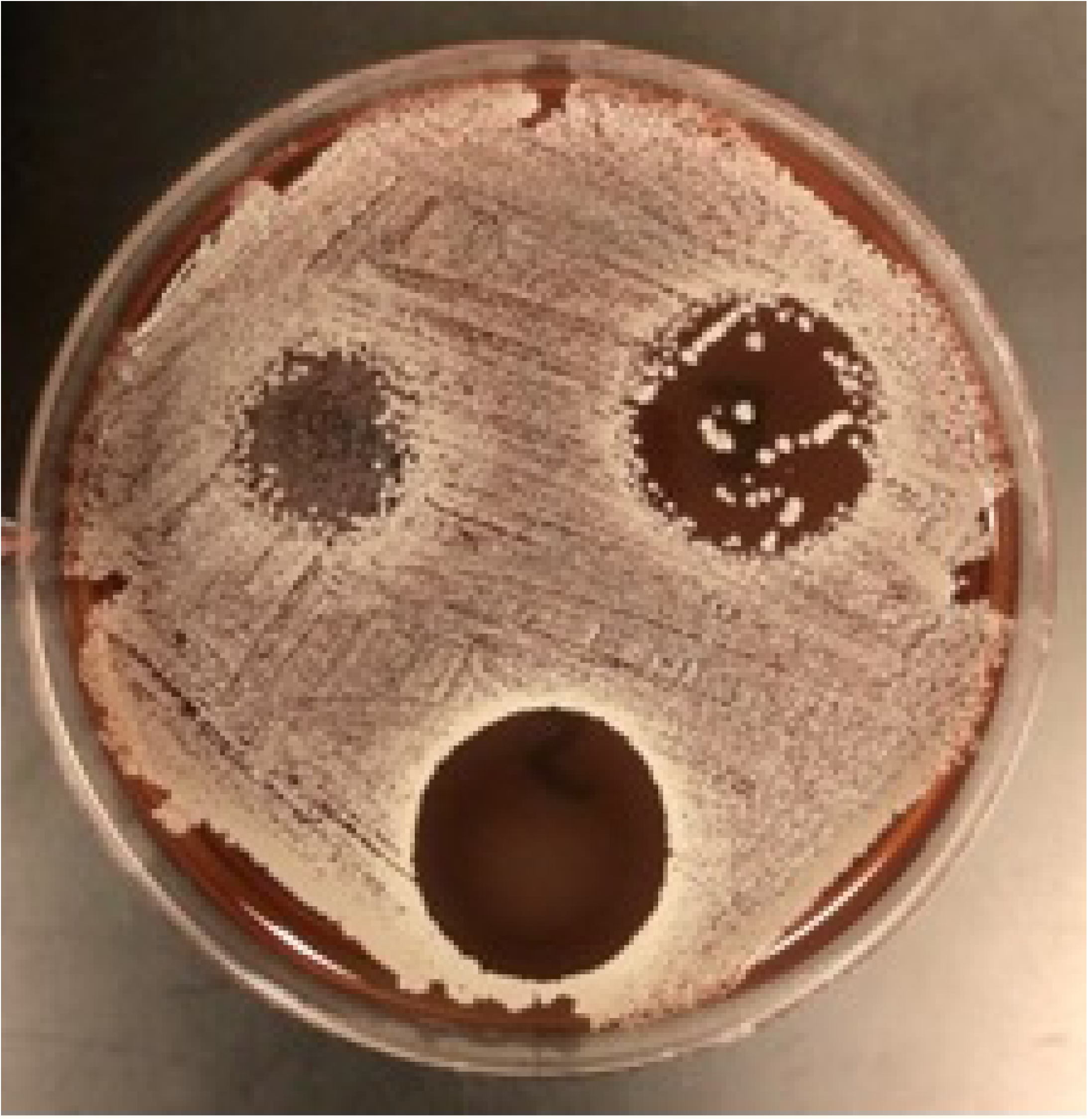
Same MRSA plate after 120 hours in incubator. Ethyl alcohol (top left), Ethanol (top right), and BAC (bottom).

**Fig 5.**
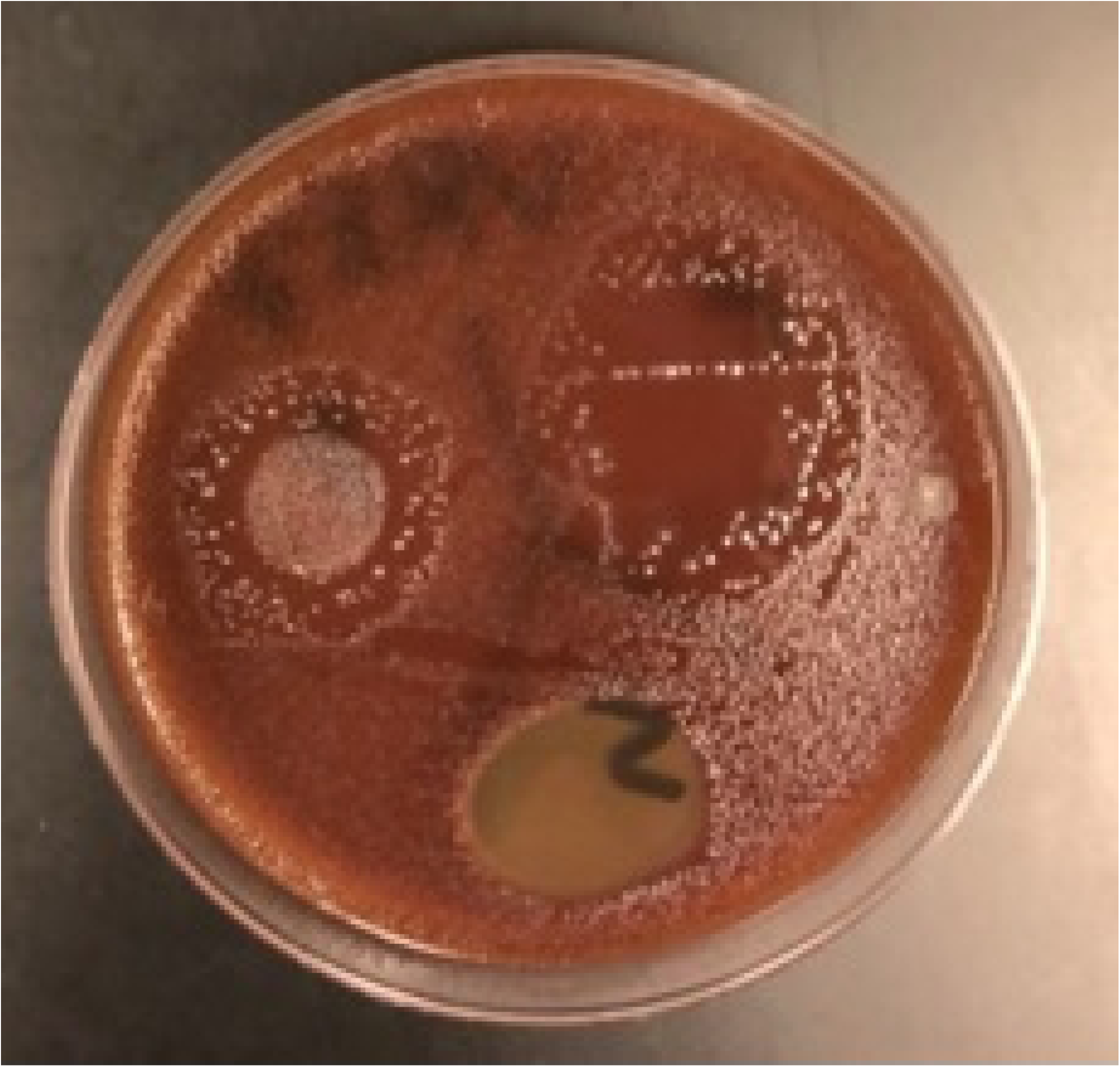
Same VRE plate after 120 hours in incubator. Ethyl alcohol (top left), Ethanol (top right), and BAC (bottom).

**Fig 6.**
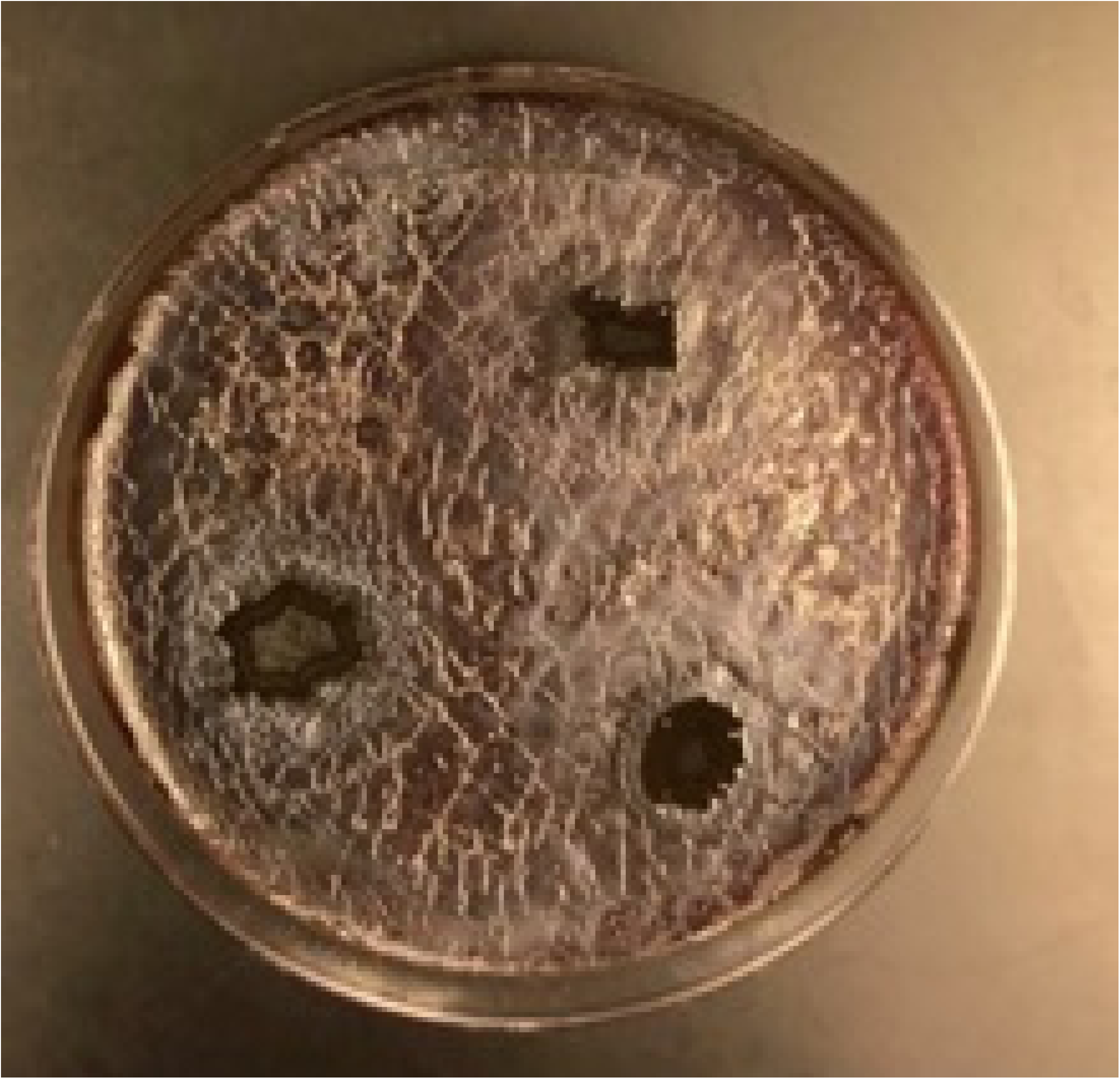
Same P. *aeruginosa* plate after 120 hours in incubator. Ethyl alcohol (top left), Ethanol (top right), and BAC (bottom).

For P. *aeruginosa*, BAC showed larger zones of inhibition on day 2, 3 and 5, but ethanol showed larger zones of inhibition on days 1 and 4. This is shown in Table 1. Aside from days 1 and 4 of pseudomonas, Table 1, Ethyl alcohol did not demonstrate a clear zone of inhibition for any other plates and thus was labeled as 0 due to significant regrowth of bacteria beginning at 24 hours and lasting the full 120 hours (Figures 1-6). Overall, BAC showed larger zones of inhibition compared to ethanol and ethyl alcohol for VRE and MRSA. This is demonstrated in Tables 2 and 3.

**Table 1.**
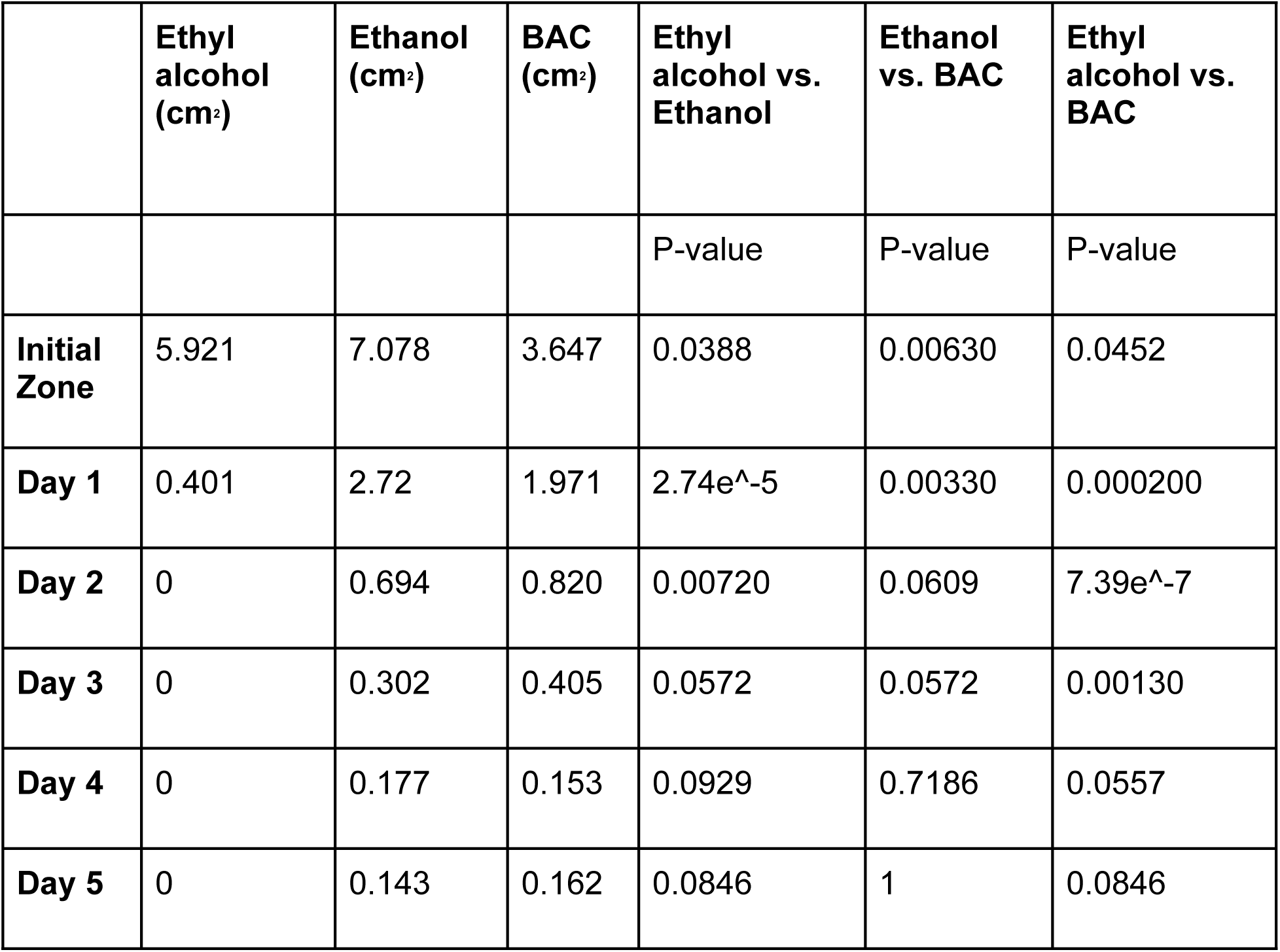
Growth of P. *aeruginosa* vs Ethyl alcohol, Ethanol, and BAC with associated p-values. Values denote the zone of inhibition measured in centimeters^2^.

**Table 2.**
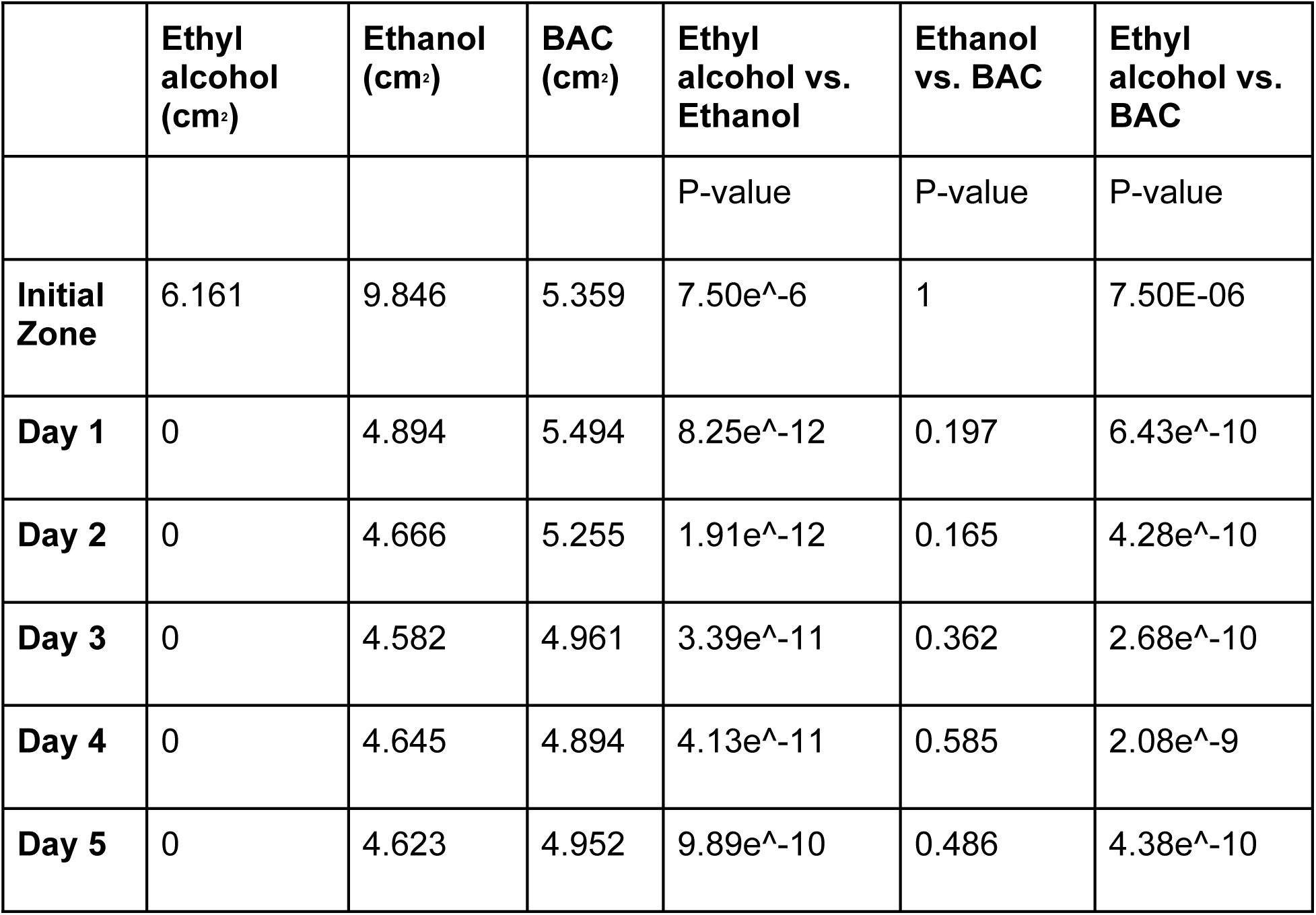
Growth of VRE vs Ethyl alcohol, Ethanol, and BAC with associated p-values. Values denote the zone of inhibition measured in centimeters^2^.

**Table 3.** Growth of MRSA vs Ethyl alcohol, Ethanol, and BAC with associated p-values. Values denote the zone of inhibition measured in centimeters^2^.

|  | **Ethyl alcohol** | **Ethanol**<br>**(cm<sup>2</sup>)** | **BAC**<br>**(cm<sup>2</sup>)** | **Ethyl alcohol**<br>**vs. Ethanol** | **Ethanol**<br>**vs. BAC** | **Ethyl alcohol vs.**<br>**BAC** |
| --- | --- | --- | --- | --- | --- | --- |
|  | **(cm<sup>2</sup>)** |  |  |  |  |  |
|  |  |  |  | P-value | P-value | P-value |
| **Initial Zone** | 1.785 | 4.035 | 5.396 | 9.56e<sup>-8</sup> | 2.41e<sup>-5</sup> | 6.49e<sup>-11</sup> |
| **Day 1** | 0 | 3.477 | 5.338 | 1.0426e<sup>-11</sup> | 3.33e<sup>-8</sup> | 6.38<sup>-17</sup> |
| **Day 2** | 0 | 3.341 | 5.289 | 5.184e<sup>-12</sup> | 4.61e<sup>-8</sup> | 6.65e<sup>-18</sup> |
| **Day 3** | 0 | 3.272 | 5.293 | 2.950e<sup>-12</sup> | 1.70e<sup>-8</sup> | 7.71e<sup>-18</sup> |
| **Day 4** | 0 | 3.27 | 5.169 | 7.286e<sup>-12</sup> | 4.73e<sup>-8</sup> | 2.01e<sup>-18</sup> |
| **Day 5** | 0 | 3.151 | 5.09 | 3.015e<sup>-11</sup> | 6.69e<sup>-8</sup> | 1.71e<sup>-18</sup> |

The ratio of percent loss of initial zone vs each subsequent zone revealed that ethyl alcohol did not inhibit growth of MRSA and VRE throughout the study period (Figs 7 and 8). BAC was most effective at preventing regrowth of all three bacteria. All three solutions had significant regrowth of P. *aeruginosa* at day 5; BAC had a regrowth of 95.5%, Ethanol 98.0% and ethyl alcohol 100% (Fig 9). For VRE by day 5, BAC showed to have a 7.6% regrowth of the initial zone vs Ethanol’s 53.0% and ethyl alcohol’s 100% regrowth (Fig 8). The day 5 regrowth of MRSA vs BAC was shown to be only 5.7% while it was 21.9% and 100% for ethanol and ethyl alcohol, respectively (Fig 7). Overall, BAC did not exhibit any regrowth within the initial zone of inhibition vs MRSA and VRE (Figs 1 and 2, 4 and 5).

**Fig 7.**
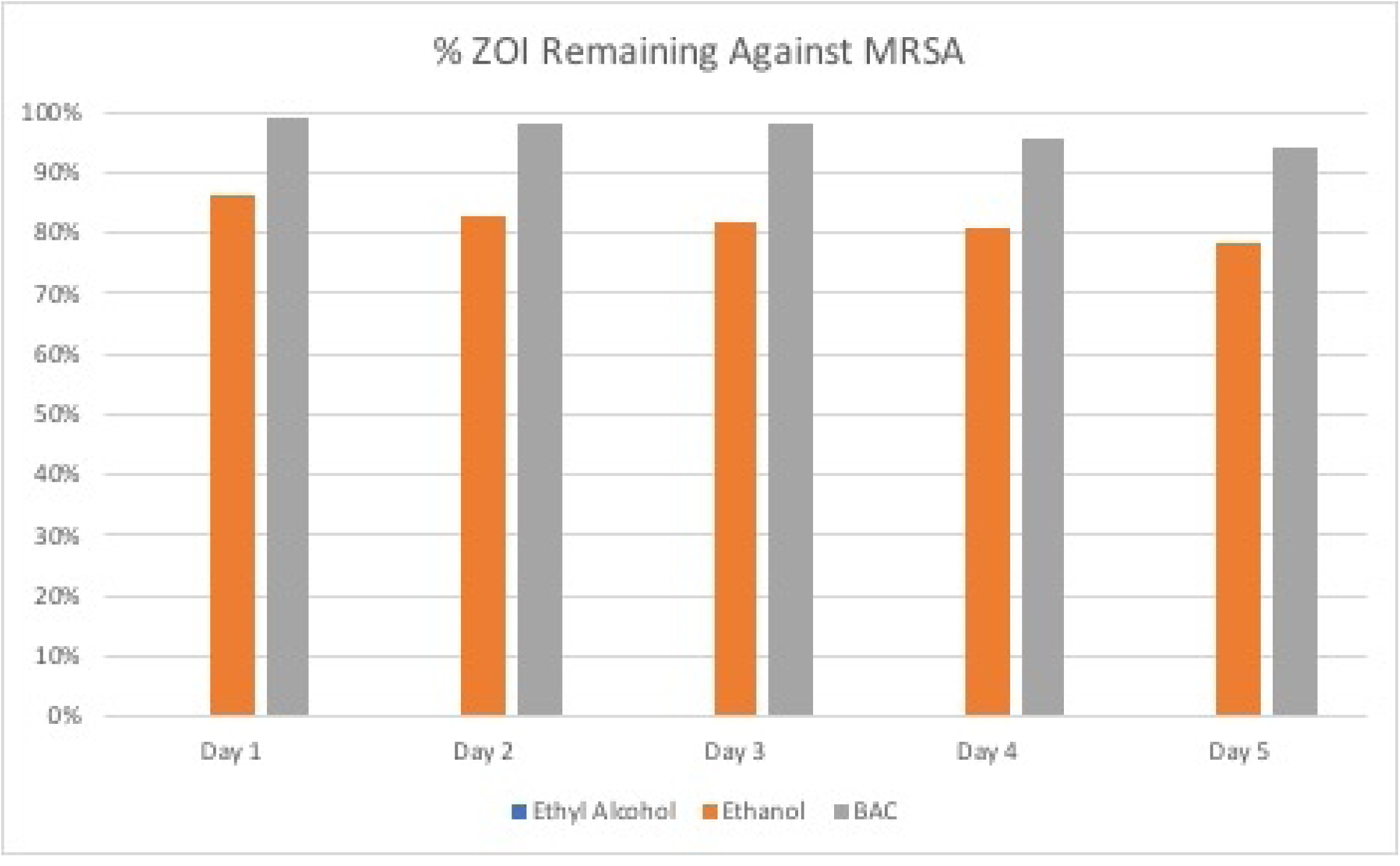
Percent of the initial zone of inhibition remaining against MRSA for each subsequent day vs Ethyl alcohol, ethanol and BAC.

**Fig 8.**
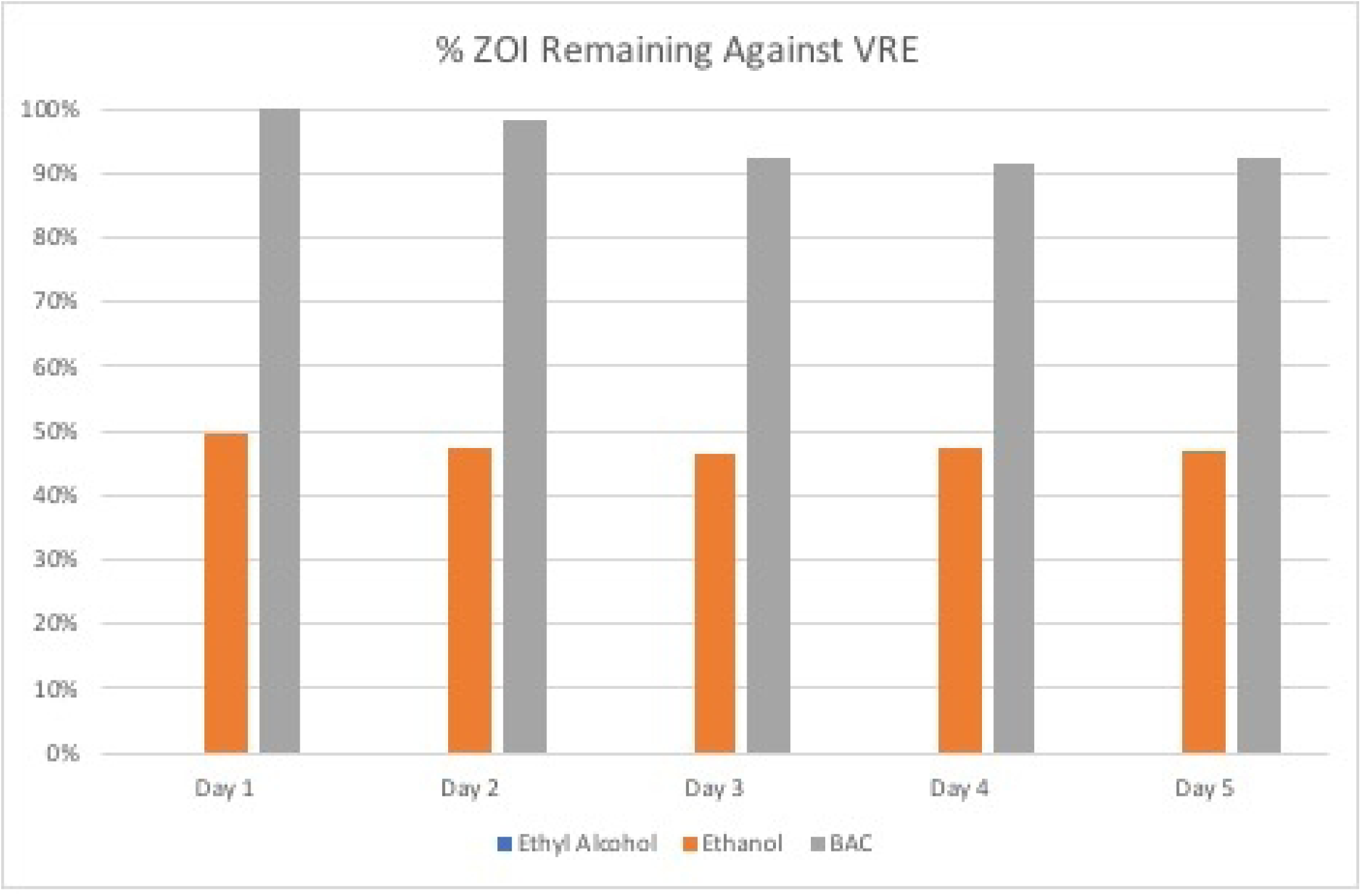
Percent of the initial zone of inhibition remaining against VRE for each subsequent day vs Ethyl alcohol, ethanol and BAC.

**Fig 9.**
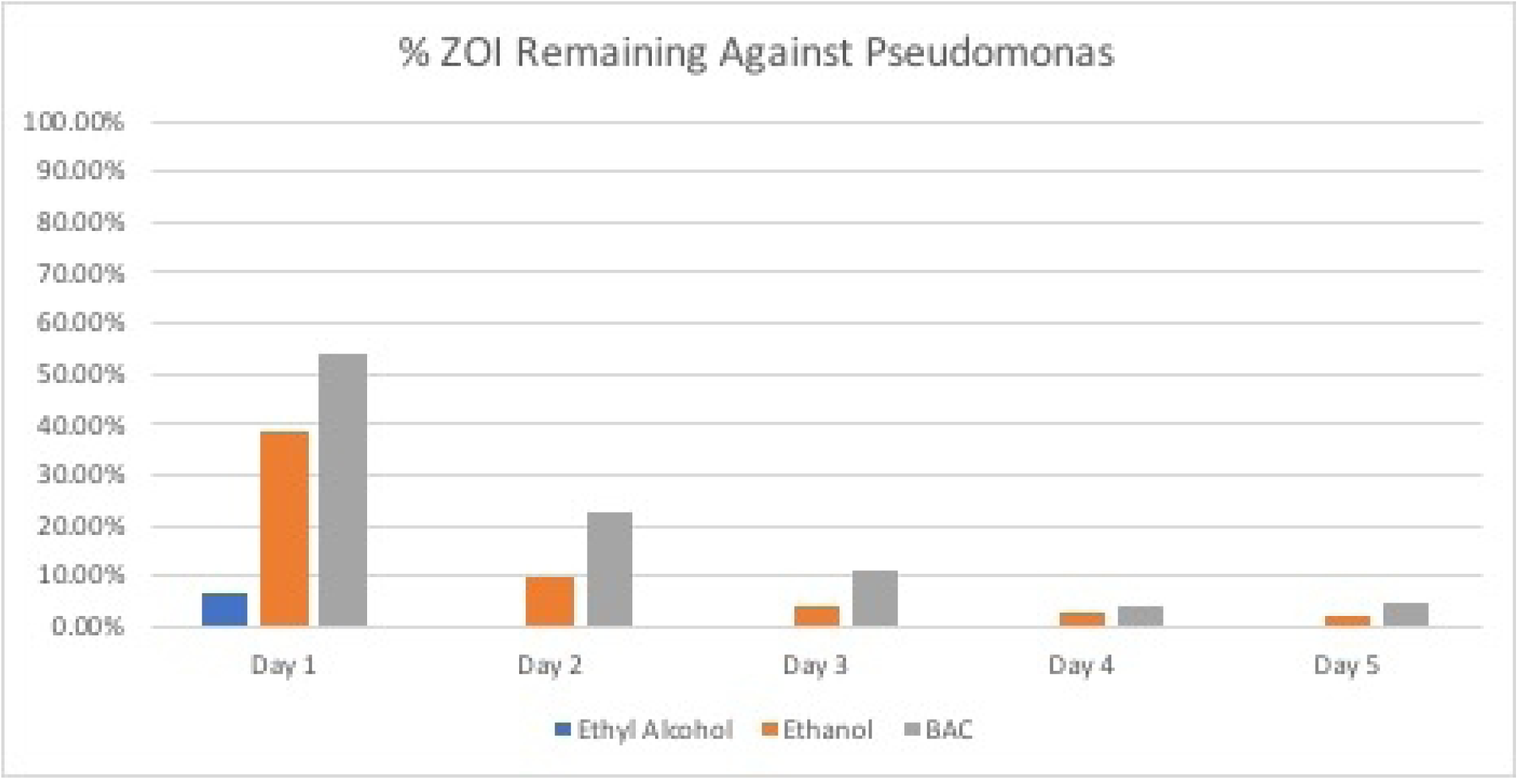
Percent of the initial zone of inhibition remaining against P. *aeruginosa* for each subsequent day vs Ethyl alcohol, ethanol and BAC.

## Discussion

Causative agents of HAIs and the routes by which they spread have been well documented for over a century. Despite knowledge of the factors that influence the risks of HAIs and means to prevent or control them [11], patients in the healthcare setting continue to acquire HAIs [8]. These infections result in increased morbidity and mortality for the patient, and ultimately higher health care costs for both the patient and the health care system [2, 12]. As one example, it’s estimated that preventing a single postoperative surgical site infection could potentially save the healthcare system upwards of $60,000 [9].

The overlying goal of this study was to test the effectiveness and duration of effect of hand sanitizers that could be used in the healthcare setting to potentially decrease the incidence of HAIs. This study demonstrates the anti-microbial activity of alcohol-based hand sanitizing agents (70% ethyl alcohol and 100% ethanol) and alcohol-free agents (BAC) against common virulent antibiotic-resistant micro-organisms MRSA, VRE, and P. *aeruginosa* in vitro over a period of 120 hours. The 0.13% BAC formulation used here exhibited superior effectiveness and duration when compared to 70% ethyl alcohol and 100% ethanol against all three micro-organisms used in this study. Specifically, against MRSA and VRE, BAC created clear cut zones of inhibition (ZOI) with minimal bacterial regrowth over a period of 120 hours. Against P. *aeruginosa*, BAC created a clear cut ZOI up to 24 hours before allowing some bacterial regrowth to occur on a few plates after the 24-hour mark, and eventually all plates after the 96-hour mark. These results are in stark contrast to the efficacy of the commonly used alcohol-based hand sanitizing agent 70% ethyl alcohol, which failed to prevent bacterial regrowth for any micro-organism. Even at the 24-hour mark no clear cut ZOI could be appreciated on any plate. Likewise, 100% ethanol also failed to prevent bacterial regrowth for any of the micro-organisms used. BAC not only proved efficacious in preventing the regrowth of bacteria over an extended period of time, but against MRSA and VRE also demonstrated an ability to maintain the integrity of the ZOI once initially established, in regard to size, over the course of 120 hours. Against P. *aeruginosa*, the ZOI created by BAC was noted to shrink in size over the duration of the experiment.

When measuring the difference in area of inhibition between each hand sanitizing agent against each bacterium, BAC proved to be most efficacious in subduing the growth of MRSA over all time points with statistically significant results (Table 3). BAC was also statistically more efficacious when compared to 70% ethyl alcohol over all time points against VRE (Table 2), and over the course of the first 72 hours against pseudomonas (Table 1). Our results showed that BAC was more efficacious against P. *aeruginosa* over the course of the first 24 hours when compared to 100% ethanol, but no significant difference was found between BAC and 100% ethanol against VRE over any time points. However, the use of 100% ethanol is unlikely as a hand sanitizing agent in the clinical setting. These results suggest that BAC could be used over 70% ethyl alcohol as a hand sanitizing agent that could provide an extended action of effective anti-microbial protection and reduce the number of HAIs spread from patient to patient by healthcare workers.

Benzalkonium chloride is commonly used in the healthcare setting as a bactericidal agent in surface disinfectants, nasal sprays, and eye drops with minimal skin irritation. The mechanism of action of BAC is related to its ability to penetrate the bacterial cell wall that leads to damage and loss of cell membrane integrity. This leads to leakage of molecular components, and eventual cell wall lysis [10]. Alcohol’s antimicrobial effect stems from its ability to denature proteins. Concentrations between 60-95% are most effective with higher concentrations losing potency because of the necessity to have water with the alcohol to be most effective [10]. Alcohol is effective at killing microbes present on the skin but has no lasting effect. BAC is non-volatile and therefore is able to remain on the skin while it dries [13]. This explains the length of duration of BACs anti-microbial effectiveness observed in our experiment. To our knowledge, the efficacy of BAC against MRSA, VRE, and P. *aeruginosa* in comparison to alcohol-based hand sanitizing agents had not been previously investigated. As such, this is the first study to demonstrate a superior ability of BAC compared to alcohol-based hand sanitizers in preventing *in vitro* MRSA, VRE, and Pseudomonas regrowth following application. Further work is necessary to determine whether BAC exhibits similar effectiveness *in vivo*.

This study has several important limitations. First, these agents were tested against microbial cultures of MRSA, VRE, and P. *aeruginosa*. It remains to be seen whether similar efficacy will be shown against these pathogens when used as a topical agent *in vivo*. Second, although the BAC formulation used in this study demonstrated a bactericidal effect the results may be variable depending on the particular strain of bacteria isolated, as modes of resistance may vary greatly between strains. Thirdly, our relatively small sample of eight bacterial plates could be the reason BAC did not show a statistical significance when compared to 100% ethanol against VRE. It remains to be seen whether a larger sample size would ultimately show that BAC is more efficacious against VRE when compared to 100% ethanol.

## Conclusion

In conclusion, BAC was shown to be superior to both 100% ethanol and 70% ethyl alcohol hand sanitizer in subduing the growth of MRSA over the course of five days. BAC was also shown to be superior when compared to 70% ethyl alcohol in subduing the growth of VRE over the course of five days, and pseudomonas over the course of 72 hours. BAC was shown to be superior to 100% ethanol in subduing the growth of P. *aeruginosa* over the course of 24 hours, but no statistically significant difference was noted over 100% ethanol against VRE. BAC was able to maintain the ZOI for VRE and MRSA throughout the course of the 5 days, whereas for P. *aeruginosa*, regrowth was observed after 24 hours. Although there was regrowth seen with BAC, it was still significantly less than the bacterial regrowth observed for 70% ethyl alcohol and 100% ethanol. The results of this study demonstrate the potential for using BAC as an effective hand sanitizer in the healthcare setting given the duration of its effect and its greater ability to potentially prevent the spread of common nosocomial infections. Further study is warranted to determine *in vivo* effectiveness of this formulation of BAC as well as the appropriate time frame of application for effectiveness against HAIs.

